# Histone methyltransferase MMSET/NSD2 is essential for generation of B1 cell compartment in mice

**DOI:** 10.1101/687806

**Authors:** Marc-Werner Dobenecker, Vyacheslav Yurchenko, Jonas Marcello, Annette Becker, Eugene Rudensky, Natarajan V. Bahnu, Thomas Carrol, Benjamin A. Garcia, Brad R. Rosenberg, Rabinder Prinjha, Alexander Tarakhovsky

**Author notes:** Abbreviations: B cell receptor (BCR), histone 3 lysine 36 di-methylation (H3K36me2), nuclear receptor SET domain-containing (NSD2). Address for correspondence: Alexander Tarakhovsky, Laboratory of Immune Cell Epigenetics and Signaling, The Rockefeller University, New York, NY 10065, USA, or Marc-Werner Dobenecker, Drug Discovery Immuno Science, Bristol-Meyers Squibb, Princeton, NJ 08540, USA.

## Abstract

Humoral immunity in mice and man relies on the function of two developmentally and functionally distinct B cell subsets - B1 and B2 cells. While B2 cells are responsible for most of the adaptive response to environmental antigens, B1 cells, which are comprised of phenotypically distinct B1a and B1b cells, are carriers of the innate humoral immunity that relies on production of poly-reactive and low affinity antibodies. The molecular mechanism of B cell specification into different subsets is not well established. Here we identified lysine methyltransferase MMSET/NSD2 as a critical regulator of the B1 cell population. We show that NSD2 deficiency in B cell precursors prevents generation of the B1 cell compartment, while having a minor impact on B2 cells. Our data revealed MMSET/NSD2, which catalyzes histone H3 lysine 36 di-methylation, as the first in class epigenetic master regulator of a major B cell lineage in mice.

## Introduction

Two developmentally and functionally distinct B cell populations support the humoral immunity in mice and man (McHeyzer-Williams, 2003). Most peripheral B cells, which are defined as B2 cells, are generated in the bone marrow and have an immensely diverse, mostly non-self directed antibody repertoire (Yurasov and Nussenzweig, 2007). B2 cells can expand rapidly upon antigen stimulation followed either by immediate differentiation into antibody producing plasma cells or by the germinal center reaction and ensuing generation of cells expressing high affinity antibody (Nutt et al., 2015). In contrast to B2 cells, B1 cells originate mostly in the yolk sack during early stages of hematopoiesis and reside predominately within well-defined anatomical compartments such as peritoneal cavity, pleural cavity and thymus (Martin and Kearney, 2001; Su and Tarakhovsky, 2000). In contrast to B2 cells, B1 cells do not proliferate in response to antigen stimulation but divide in a seemingly autonomous fashion at a low rate (Deenen and Kroese, 1992). The antibody repertoire of B1 cells is represented largely by self-reactive or poly-reactive antibody of low affinity, mostly of the IgM or IgG3 isotypes and IgA in mucosal surfaces (Baumgarth, 2011; Montecino-Rodriguez and Dorshkind, 2006; Rothstein, 1990). In addition to the markedly different developmental and functional features, B1 cells display a distinct pattern of surface proteins, including CD5 or CD11b that are expressed normally on T cells or macrophages, respectively (Forster et al., 1991; Ghosn et al., 2008; Su and Tarakhovsky, 2000). Accordingly, the CD5-positive B1 cells are defined as B1a cells and the CD5-negative B1 cells as B1b cells.

The origin of B1 cells is not well established and opposing ideas have been put forward in the literature. Some findings suggest they develop from a distinctive, and predominantly fetal lineage (Hardy and Hayakawa, 1991; Hayakawa et al., 1985), while others indicate that their phenotype is a result of the signals derived from their surface antigen receptors (Haughton et al., 1993). These concepts may not be mutually exclusive since B1 cells expresses poly-reactive antigen receptors (Kantor et al., 1997) that could be particularly amenable to stimulation from self or environmental antigens, leading to the surface expression of characteristic markers. The discovery of B1 restricted progenitors (Montecino-Rodriguez and Dorshkind, 2006; Tung et al., 2006) and of lin28b as a key regulator of B1 cell development (Yuan et al., 2012) might support the existence of a separate lineage for these cells.

Here we present data that show the essential and selective role of histone methyltransferase MMSET/NSD2 in generation of the B1 cell population in mice.

## Results and discussion

### B cell specific inactivation of MMSET/NSD2

MMSET/NSD2 (hereafter NSD2, encoded by the Whsc1 gene) is one of three members of the NSD (nuclear receptor SET domain-containing) family of histone lysine methyltransferases that contain in addition to the catalytic SET domain, PHD (plant homeodomain) fingers, PWWP (Pro-Trp-Trp-Pro motif) domains, and a NSD specific C5HCH domain (a cysteine-rich domain).

The substrate specificity of NSD2, while most likely targeting the lysine 36 of histone H3, remains controversial and largely based on *in vitro* enzymatic assays. The methylation of lysine 36 of histone H3 has been implicated in the process of RNA elongation during transcription thus suggesting NSD2 contribution to the generation of full-length transcripts (Greer and Shi, 2012). NSD2 function is essential for normal development in mice and humans and deficiency in NSD2 in mice leads to death of newborn mice due to severe growth retardation (Nimura et al., 2009). NSD2 is often deleted in Wolf-Hirschhorn syndrome and a great deal of attention for NSD2 stems from its link to aggressive myelomas in human. The t(4;14) translocation places the *Whsc1* gene under the control of the IgH Eμ-enhancer and leads to NSD2 over-expression, which is a sign of aggressive myeloma with a poor prognosis (Keats et al., 2005). The mechanism of NSD2 contribution to myelomagenesis and/or tumor progression is not understood.

With the aim to define the mechanism of NSD2 contribution to B cell or plasma cell malignancies, we first addressed the role of NSD2 in B cell generation and activation. To exclude NSD2 ability to mediate histone H3 methylation as well as methylation of other potential non-histone substrates, we have generated mice that allow Cre recombinase-mediated conditional inactivation of the catalytic SET domain of the NSD2 protein in B lineage cells (Figure 1) or other cell types. To delete the SET domain, exons 18 and 19 of the *Whsc1* gene were flanked with *lox*P sites that upon Cre mediated recombination generate a mutant *Whsc1* locus encoding a SET-domain deficient NSD2 protein. To achieve B lineage specific NSD2-SET inactivation we employed Mb1-Cre mice that allow conditional inactivation of genes starting at the pro-B cell stage of B cell development (Hobeika et al., 2006). Mb1-Cre mice were bred to *Whsc1*flSET mice and the resulting Mb1-cre; *Whsc1*flSET mice and the relevant littermate controls were used in our studies. We also generated Vav1-cre; *Whsc1*flSET mice that allow NSD2 inactivation during early stages of definitive hematopoiesis in the bone marrow (de Boer et al., 2003).

**Figure 1:**
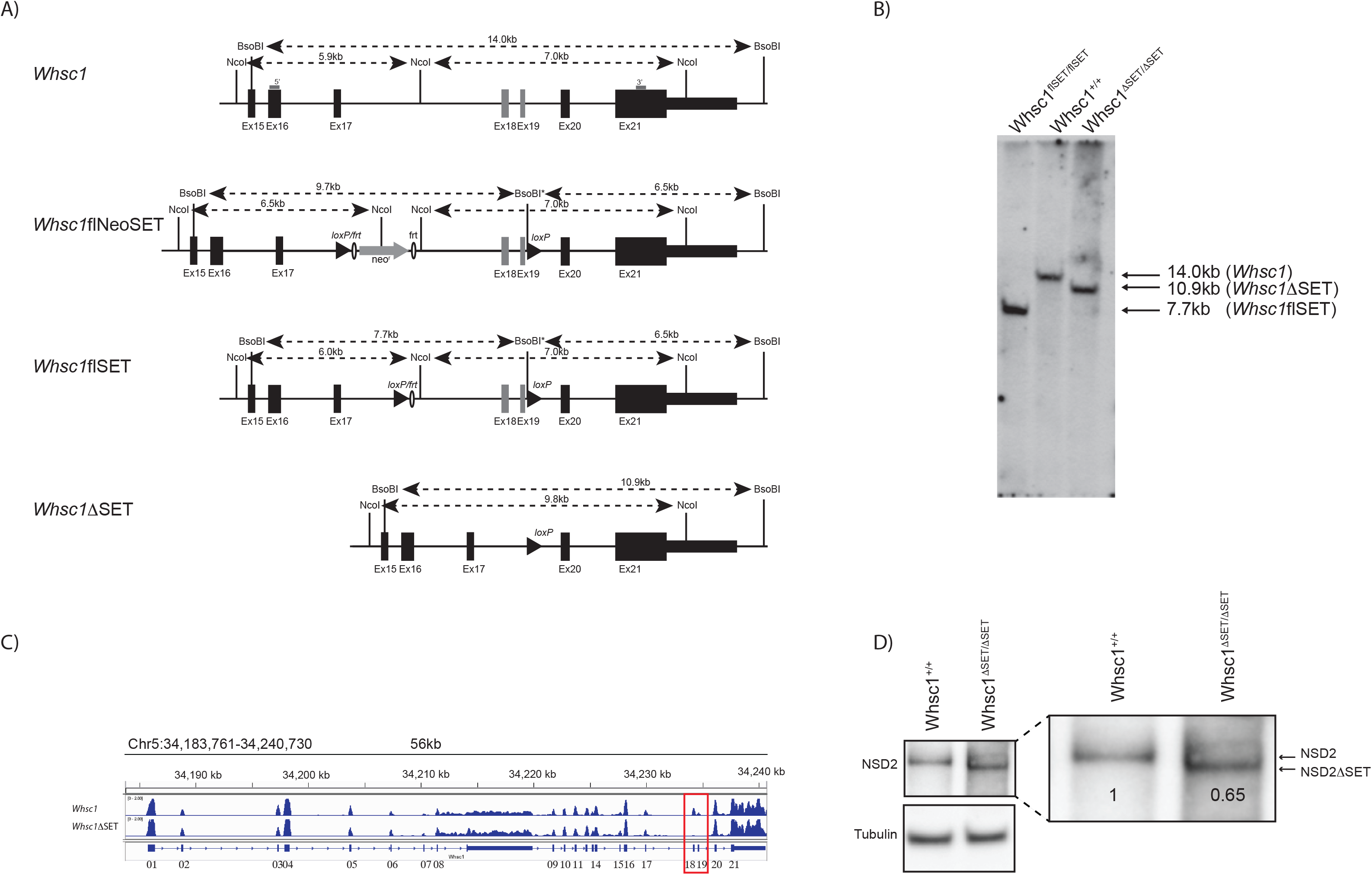
Generation of mice with B cell specific expression of catalytically inactive NSD2 (NSD2ΔSET). A) Structure of the wild type *Whsc1* locus, the targeted locus (*Whsc1*flNeoSET), the targeted locus after FLP-recombination mediated deletion of the *neo*^*r*^-cassette (*Whsc1*flSET), and the SET-domain deleted locus after Cre-recombination (*Whsc1*ΔSET). Numbered rectangles represent exons, filled triangles represent *loxP* sites, open ovals *FRT* sites; the grey arrow indicates the position and transcriptional orientation of the *neo*^*r*^ selection cassette. Restriction sites and distances are indicated above each locus. The 5’ and 3’ probes used in Southern blots are shown as grey bars. B) Mb-1 Cre mediated deletion of the SET-domain encoding exons 18 and 19 of the *Whsc1* gene in splenic B cells. Southern blot analysis of DNA isolated from purified B cell with the 5’ probe after *Bso*BI digest is shown. The 7.7 kb band corresponds to the targeted locus and the 14 kb and the 10.9 kb bands correspond to the wild-type and Cre modified *Whsc1* gene respectively. C) RNA sequencing shows the presence of the full length or SET – “less” (minus exons 18 and 19) mRNA isolated from purified B cells derived from control or Mb1-cre; *Whsc1*flSET mice, respectively. IGV tracks show the relative RNA expression level. Exons 18 and 19 encoding the SET domain are highlighted. D) Truncated NSD2ΔSET protein expression in purified B cells. The arrows indicate the wild type and truncated NSD2ΔSET protein expressed in purified B cells derived from *Whsc1*flSET or Mb1-cre; *Whsc1*flSET mice. Exons 18 and 19 encode for 83 amino acids, which are missing in NSD2ΔSET.

We found that the Cre-driven modification of *Whsc1* gene in Mb1-cre; *Whsc1*flSET mice leads to the expression of a truncated SET domain-deficient NSD2 protein (hereafter NSD2ΔSET). The expression levels of the mutant NSD2 protein levels are approximately 30% lower than the full length NSD2 protein in wild-type B cell (Figure 1d). The expression of NSD2ΔSET in B cells results in selective changes in the pattern of histone H3 post-translational modification in B2 cells. In agreement with the previously reported NSD2 specificity towards di-methylation of lysine 36 of histone H3 (H3K36me2) (Li et al., 2009), we found greatly reduced levels of H3K36me2 in B cells. Histones extracted from wild type and NSD2ΔSET splenic B2 cells were analyzed by mass-spectrometry for the presence of various histone modifications. The histones isolated from NSD2ΔSET B cells showed only few significant changes, mainly in H3K36 methylation in addition to other modifications, which we deem secondary. By far most significant are major and selective reductions of H3K36me2 and H3K36me3 and a corresponding increase in unmodified H3K36 as compared to the histones derived from control cells (Figure 2A, B).

**Figure 2:**
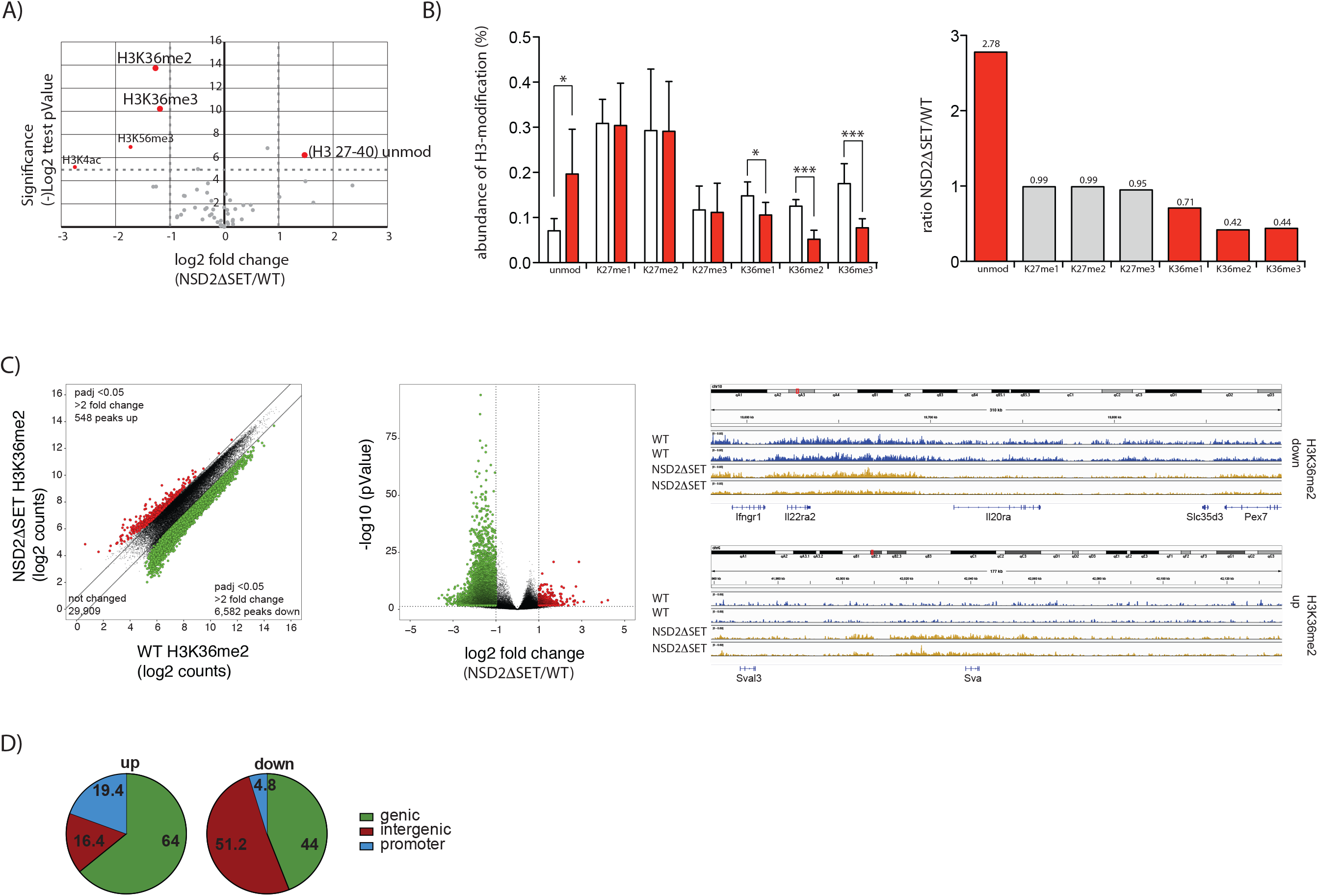
Selective methylation of histone H3 at lysine 36 by NSD2. A) Volcano plot shows changes in relative abundance of distinct modifications of histones extracted from splenic B cells purified from wild-type or mutant mice. Values over 4.32 in the Y-axis [Log2 (NSD2ΔSET/WT)] corresponding to -log2 (0.05) are significant and highlighted in red. The data represents two biological replicates and three independent measurements. Significance was determined by unpaired Student’s T-test. *: p ≤ 0.05, **: p ≤ 0.01, ***: p ≤ 0.001. B) The fraction of histone H3 modified at lysine 36 was determined by combining the frequencies for all histone H3 peptides carrying this modification, independent of other modifications present on the same peptide. Peptides of controls are represented by white bars, peptides from Mb1-cre; *Whsc1*flSET mice in red bars (left). Fold change of H3-peptides carrying the indicated modifications. The significant changes in abundance are indicated in red bars, not significant changes in grey bars. Numbers above the bars indicate the fold change of mutant over wild-type (right). C) Scatter plot of H3K36me2 peaks of splenic B cells from WT and Mb1-cre; *Whsc1*flSET mice. Significant increase (red) or decrease (green) in specific peaks are indicated (left). Volcano plot illustrating differential H3K36me2 levels in splenic B cells from WT and Mb1-cre; *Whsc1*flSET mice. The log2 fold change in abundance (NSD2ΔSET/WT) is plotted along the X-axis and the -log10 pValue is plotted on the Y-axis (p=0.05 corresponds to 4.32). Significant increase (red) or decrease (green) in specific peaks are indicated (right). Data plotted are average normalized H3K36me2 values of called peaks from three biological replicates.

The reduction in H3K36me2/3 methylation affects over 50% of all thus modified histones (Figure 2B) and the loss of H3K36me2/3 is independent of other modifications on the same histone (data not shown). This result strongly supports the H3K36me2 specificity of NSD2. ChIP-seq data for H3K36me2 shows a locus specific and significant (p-adjusted <0.05) reduction of over two-fold at 6,582 peaks and an increase of over two-fold at 548 peaks (Figure 2C). Sample IGV tracks confirm the locus specific changes that occur on the scale of serval to hundreds of kb (Figure 2C). The gain of H3K36me2 in NSD2ΔSET B cells occurred mainly in genic (64%) and promoter regions (19.4%) while the loss of H3K36me2 was observed mainly in intergenic regions (51.2%) followed by genic regions (44%) (Figure 2D). The selectivity of H3K36me2 down-regulation only at some gene targets suggest a locus-specific mechanism of NSD2 targeting to the chromatin in B2 cells. How such specificity is achieved remains to be understood. A likely scenario is that NSD2 is recruited to chromatin with the help of its non-catalytic domains, which differ between distinct NSD family members (Wagner and Carpenter, 2012).

Furthermore, the inactivation of the SET domain in NSD2 reproduces the effect of *Whsc1* gene null-mutation. We generated mice with germ-line inactivation of SET-NSD2 followed by breeding of NSD2ΔSET to homozygosity. We found that similar to NSD2-deficient mice (Nimura et al., 2009), NSD2ΔSET homozygous mice die early after birth. This result allows us to relate the NSD2ΔSET mutation to NSD2 deficiency.

### NSD2 is dispensable for B2 cell development and activation

The wild-type like pattern of B cell development in the bone marrow of Mb1-cre; *Whsc1*flSET mice shows that NSD2 is not essential for B2 cell development (Figure 3A). The unaltered pro-B to pre-B cell transition argues against a critical role of NSD2 in IgH gene rearrangement and expression as well as division of B cell progenitors. The wild-type like pattern of immature B2 cell generation in the bone marrow of Mb1-cre; *Whsc1*flSET mice also points to a lack of NSD2 significance in processes required for surface expression of IgM and signaling required for the generation of immature B cells. The number of re-circulating cells in the bone marrow and of follicular B2 cells in the spleen are reduced (Figure 3B). NSD2 might therefore play a role in the regulation of peripheral B2 cell maintenance, including B2 cell division or survival. We tested the impact of NSD2ΔSET on proliferation in response to antigen receptor-or polyclonaly-triggered B cell proliferation *in vitro* and found no defect (Figure 3C). Also, the ability of B2 cell to respond to pro-survival signals such as BAFF was not affected by NSD2-SET deficiency (Figure 3D). Incubation of the mutant B2 cells with BAFF supports *in vitro* cell survival in a fashion similar to wild-type B cells. Overall, our data show a non-essential role of NSD2 in generation of the B2 cell compartment. One of the major conclusions of these studies is related to the obvious lack of NSD2 contribution to B cell division. This particular finding argues against the current view on NSD2 as an important regulator of cell proliferation.

**Figure 3:**
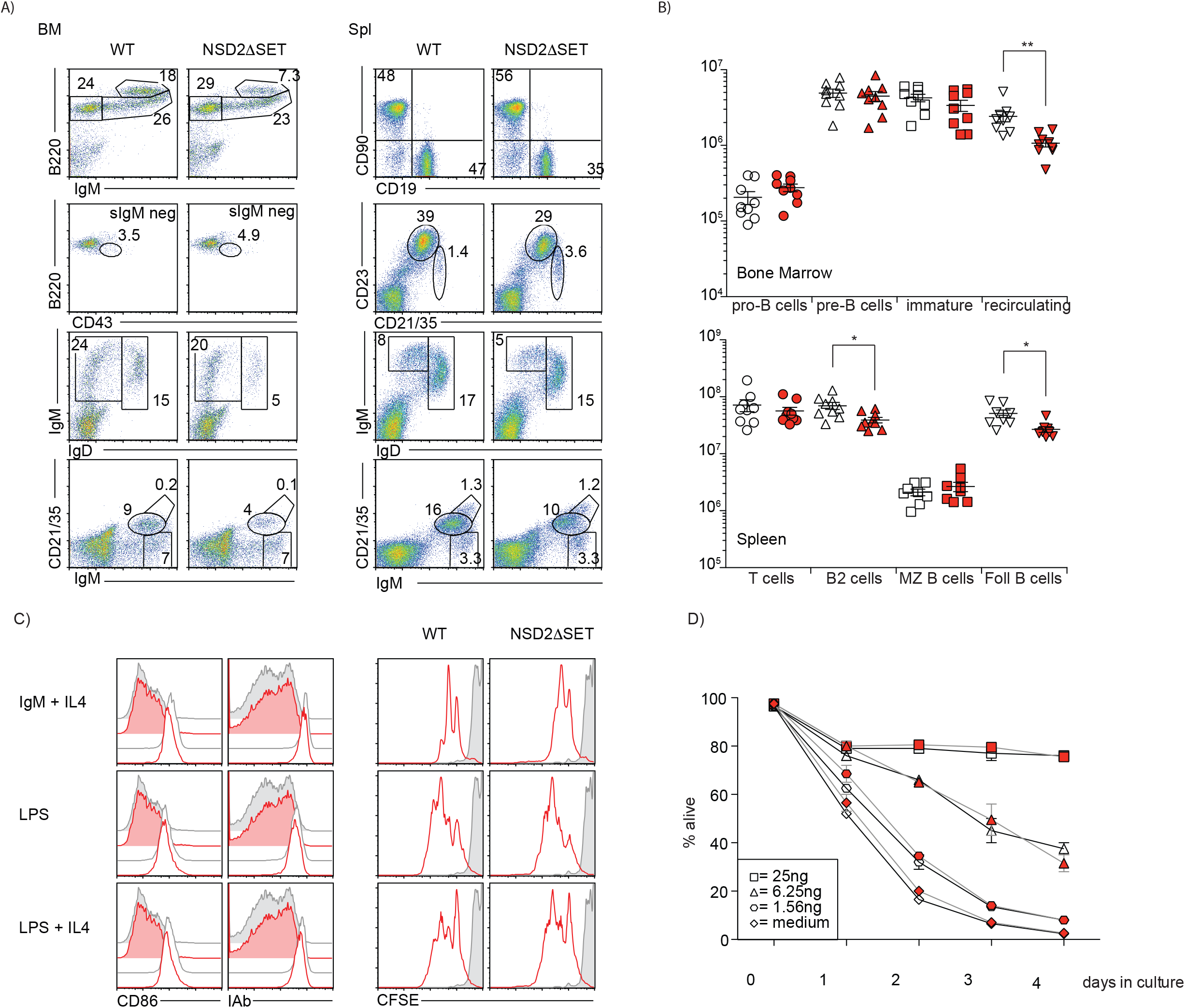
NSD2 is not essential for the development and maturation of B2 cells. A) FACS dot plots show distribution of developing and mature peripheral B2 cells. The percentage of cells of defined phenotypes are indicated. Representative plots of more than 3 independent experiments with 3 or more mice per group are shown. B) The absolute numbers of developing and mature B2 cells in the bone marrow (top) and spleen (bottom) of *Whsc1*flSET control (white symbols) and Mb1-cre; *Whsc1*flSET (red symbols) mice are shown. Each symbol represents one mouse. The experiments were performed 3-4 times with 2-3 mice per genotype. Significance was determined by unpaired T-test *: p ≤ 0.05, **: p ≤ 0.01. C) Dispensable role of NSD2 catalytic function in B cell proliferation. Expression of CD86 and IA^b^ on freshly isolated WT (grey) and NSD2ΔSET (red) B cells in the absence (filled histograms) or presence of stimuli (open histograms). Proliferation of freshly isolated WT and NSD2ΔSET B cells in the absence (filled histograms) or presence of stimuli (open histograms) was measured by CFSE dilution. The experiments were performed 3-4 times with 2-3 mice per genotype. D) Dispensable role of NSD2 catalytic function in B cell survival. Frequencies of viable WT (open symbols) and NSD2ΔSET (red symbols) B cells in the presence of 0-25 ng recombinant Baff. Data are means ± s.d. from 3 independent experiments.

### NSD2 controls switching to IgG3 and IgA

Previous studies reported that NSD2 was required for B cell class switch recombination (Nguyen et al., 2017; Chen et al., 2018) and in accordance with these studies we also found reduced serum Ig levels of IgM, IgG3, and IgA (Figure 4A). Mb1-cre; *Whsc1*flSET mice have a reduced percentage of IgA positive B cells in Peyer’s Patches (Figure 4B) and NSD2ΔSET B cells have a reduced ability to switch to IgA *in vitro* (Figure 4C). Under all tested conditions, with varying amounts of TGF-β or All Trans Retinoic Acid (ATRA) NSD2ΔSET B cells are always less efficient in switching to IgA, while they maintained their ability to upregulate the integrin α4β7, a known target of RA-signaling (Iwata et al., 2004). Similarly, switching to IgG3 in response to LPS is also impaired, while switching to IgG1 in response to LPS+IL4 showed no defect (Figure 4D). Apart from the switching defect we also found that the splenic germinal center response was reduced by ∼50% in Mb1-cre; *Whsc1*flSET mice in response to the model antigen Sheep Red Blood Cells (SRBCs) but maintained the characteristic increase of cells in the light zone (LZ) (Figure 4E). Immunization with the T-independent model antigen NP-Ficoll resulted in the characteristic induction of antigen specific IgM, IgG3, and λ-chain carrying antibodies, with a slight reduction in the number of IgM specific antibodies at day 14 post immunization (Figure 4F). In response to immunization with the T-cell dependent antigen NP_22_-CGG we found a defect in the recall response to the secondary immunization for IgG1 and a general defect in the induction of antigen specific antibodies of the IgG3 isotype (Figure 4G). These findings argue for a role of NSD2 in isotype switching that surprisingly is restricted to a limited number of isotypes. How this specificity is achieved is currently unknown.

**Figure 4:**
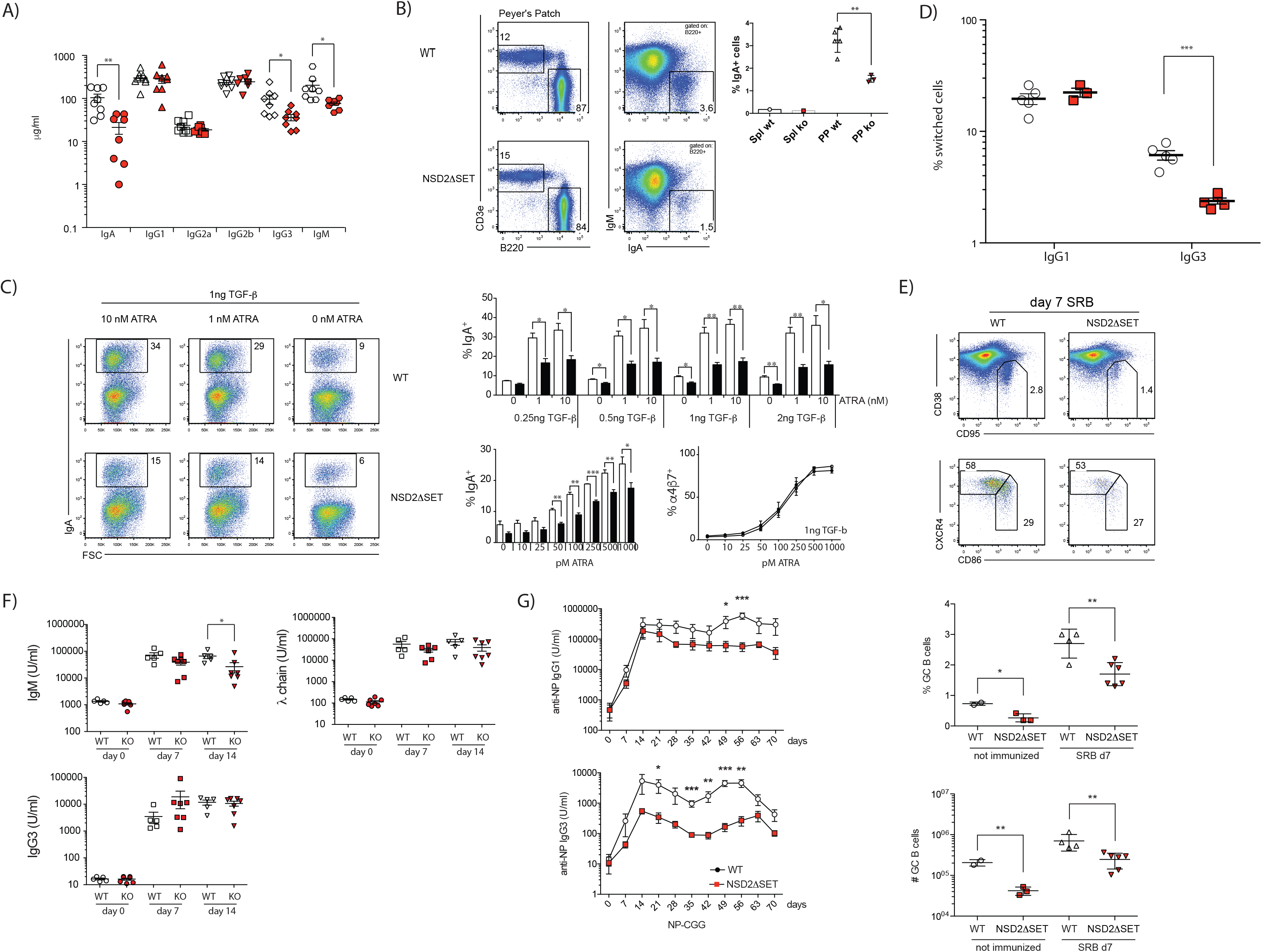
NSD2 catalytic function controls selective class switch recombination in B cell. A) Concentration of serum immunoglobulins in control (white symbols) and Mb1-cre; *Whsc1*flSET mice (red symbols) were quantified by ELISA. Each symbol represents one mouse. Significance was determined by unpaired T-test *: p ≤ 0.05, **: p ≤ 0.01, ***: p ≤ 0.001. B) FACS plots show relative abundance of T cells (CD3e^+^) and B2 cells (B220^+^); and the relative abundance of surface IgA positive (B220^+^IgA^+^) B cells in Peyer’s Patches of control and Mb1-cre; *Whsc1*flSET mice (NSD2ΔSET). The percent of IgA^+^ B2 cells in spleen (Spl) and Peyer’s Patches (PP) is indicated on the right. C) NSD2ΔSET B2 cells have a defect in switching to IgA *in vitro*. B2 cells isolated from control and Mb1-cre; *Whsc1*flSET mice were stimulated *in vitro* in the presence of LPS, TGFβ, and all trans retinoic acid (ATRA). The percentage of IgA positive B cells is indicated. Representative plots of more than 3 independent experiments with 3 or more mice per group are shown. Bar diagrams indicate the percent of IgA positive B2 cells after 3 days in various culture conditions. Control cells (white bars) and cells from Mb1-cre; *Whsc1*flSET (black bars) mice (top). NSD2ΔSET B2 cells have a defect in switching to IgA but not in the induction of the integrin α4β7 in response to ATRA. The percentage of IgA positive B cells is response to LPS and a stable amount of TGFβ (1ng) with increasing amounts of ATRA (0-1000pM) in control (white bars) and NSD2ΔSET (black bars) B2 cells is indicated (bottom left). The percentage of integrin α4β7 positive B cells is response to LPS and a stable amount of TGFβ (1ng) with increasing amounts of ATRA (0-1000pM) in control (open symbols) and NSD2ΔSET B2 cells (closed symbols) is indicated (bottom right). D) NSD2ΔSET B2 cells have a defect in switching to IgG3 but not IgG1*in vitro*. B2 cells isolated from control (open symbols) and Mb1-cre; *Whsc1*flSET (red symbols) mice were stimulated *in vitro* in the presence of LPS or LPS+IL4. The percentages of switched B cells after 3 days *in vitro* culture are indicated. Each symbol represents one mouse. Significance was determined by unpaired T-test *: p ≤ 0.05, **: p ≤ 0.01, ***: p ≤ 0.001. E) B cells from Mb1-cre; *Whsc1*flSET (NSD2ΔSET) mice have a defect in entering the germinal center reaction. Control and Mb1-cre; *Whsc1*flSET (NSD2ΔSET) mice were injected IP with sheep red blood cells and the percent of germinal center (GC) cells (CD95^+^CD38^dull^) and the percent of GC B cells in the light zone (CD86^+^CXCR4^dull^) and dark zone (CD86^−^CXCR4^+^) are indicated in the representative FACS plots. The percent and total number of GC B cells in control (open symbols) and Mb1-cre; *Whsc1*flSET (NSD2ΔSET) mice (red symbols) is plotted below. Each symbol represents one mouse. Significance was determined by unpaired T-test *: p ≤ 0.05, **: p ≤ 0.01. F) T independent immunization. Control (WT, open symbols) and Mb1-cre; *Whsc1*flSET (KO, red symbols) mice were immunized with the T independent model antigen NP-Ficoll and titers of the antigen specific antibodies were determined by ELISA. G) T dependent immunization. Control (WT, open symbols) and Mb1-cre; *Whsc1*flSET (KO, red symbols) mice were immunized (day 0) and boosted (day 35) with the T dependent model antigen NP_22_-CGG and titers of the antigen specific antibodies were determined by ELISA. Each symbol represents one mouse. Significance was determined by unpaired T-test *: p ≤ 0.05, **: p ≤ 0.01, ***: p ≤ 0.001. The experiment was performed once with 5-8 mice per experimental group.

### NSD2 is essential for generation of B1 cell compartment

Contrary to B2 cells, peritoneal B1 cells were absent in Mb1-cre; *Whsc1*flSET mice. FACS analysis of peritoneal cells derived from Mb1-cre; *Whsc1*flSET mice show an over 12-fold reduction in the number of IgM^+^CD5^+^CD11b^+^ B1a cells and a nearly 3-fold reduction of the B1b cells (Figure 5A, B). The major reduction of B1 cells numbers is mirrored by changes in serum antibody titers (Figure 4A). It is well established that B1 cells are the largest contributors to the overall serum levels of IgM and IgG3, which in addition to the switching defect in B2 cells (Figure 4C, D), can explain the reduction in serum levels of these Igs. Notably the CD5^hi^ and phosphotidylcholine-specific BCR expressing cells are completely absent in Mb1-cre; *Whsc1*flSET mice (Figure 5A). The phosphotidylcholine specificity is determined largely by Vh11/Vh12 bearing heavy chains paired with various lights chains, most favorably Vk4 or Vk9 (Clarke and McCray, 1993). Therefore, lack of phosphotidylcholine-specific B cells (Figure 5C) may potentially reflect defective Vh11/Vh12 chain rearrangement and/or expression. Analysis of NSD2 expression in B1a, B1b, and B cells of Mb1-cre; *Whsc1*flSET mice revealed that the residual B1 cells still express the unmodified wild-type NSD2 transcript, and were therefore not deleted for the SET domain, whereas B2 cells express only the NSD2ΔSET transcript (Figure 5D). To address the question whether NSD2 is essential for the generation or the maintenance of B1 cells we analyzed 3-week old mice. While littermate control mice had a large percentage of B1a and B1b cells, young Mb1-cre; *Whsc1*flSET mice displayed a significant reduction in the percent and number of these populations, indicating that the development and not the maintenance of B1 cells is impaired. Recently the role of lin28 in fetal haematopoiesis was described (Yuan et al., 2012). Ectopic expression of lin28b results in increased development of B1a, marginal zone (MZ) B, gamma/delta (γδ) T cells, and natural killer T cells. NSD2ΔSET only effects the development of B1 and natural killer T cells but not of MZ B cells or γδ T cells (Figure 3, 5, and data not shown), making it unlikely that the defect in B1 cell development is the consequence of changed lin28 expression in NSD2ΔSET cells.

**Figure 5:**
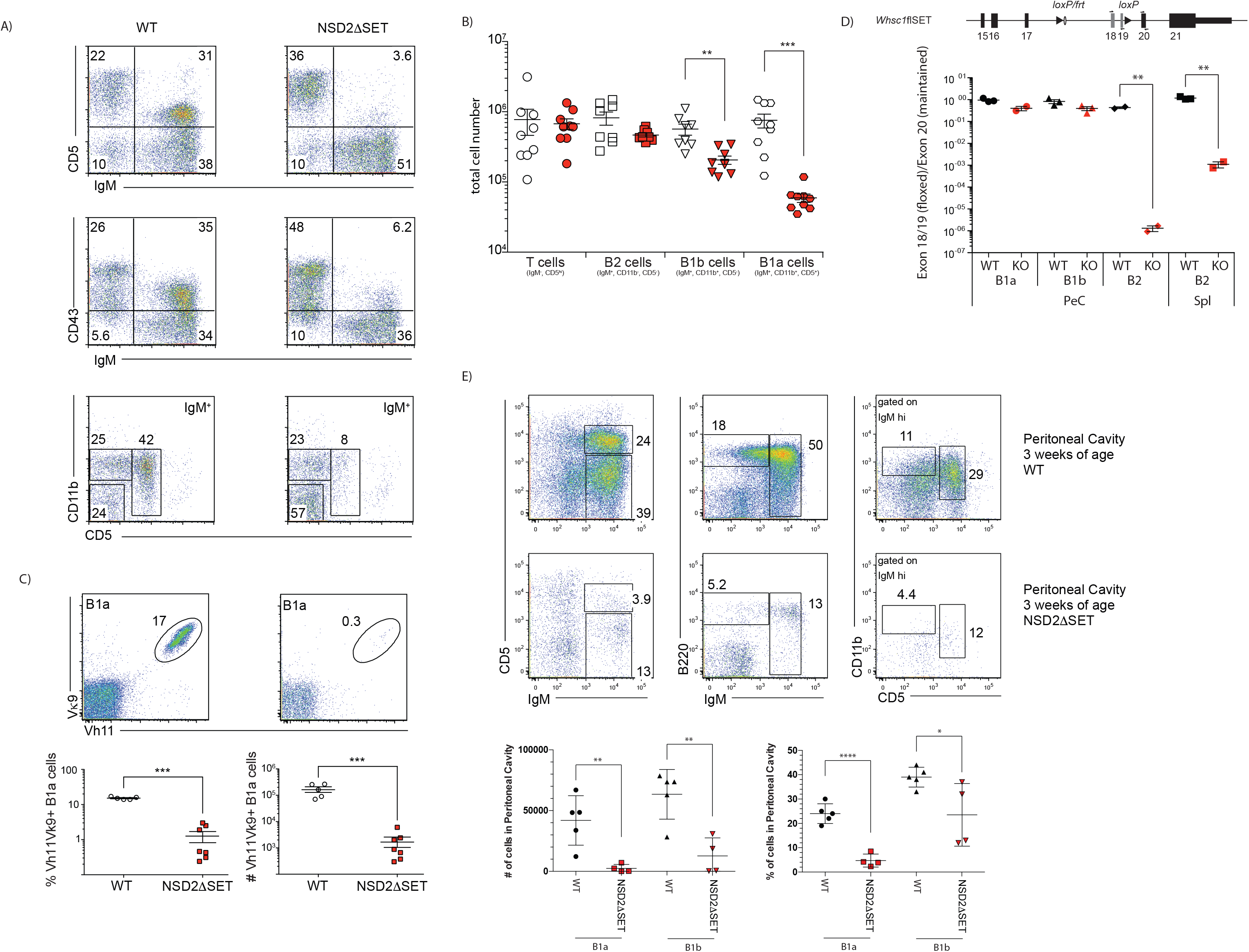
NSD2 is essential for B1 cell generation. A) FACS plots show relative abundance of T cells (CD5^+^IgM^−^), B1a (IgM^hi^CD11b^+^CD5^+^); and B1b (IgM^hi^CD11b^+^ CD5^−^) and B2 (IgM^+^CD11b^−^CD5-) cells. B) Absolute numbers of distinct lymphoid cells in the peritoneal cavity. Representative plots of more than 3 independent experiments with 3 or more mice per group are shown. C) The frequency (bottom left) and absolute number (bottom right) of B1a cells expressing phosphatidylcholine specific B cells receptor (Vκ9 Vh11) in controls (white symbols) or Mb1-cre; *Whsc1*flSET mice (red symbols) are shown. Representative plots of 2 independent experiments with six mice in total are shown. D) Deletion efficiency was estimated by calculating the ratio of qPCR products for exons 18/19 (flanked by *loxP* sites) over Exon 20 (maintained). Numbered rectangles represent exons, filled triangles represent *loxP* sites, and arrows indicate pPCR primers used to amplify either exons 18+19 or exon 20 (top). The ratio of the qPCR products from control cells (black symbols) or cells from Mb1-cre; *Whsc1*flSET mice (red symbols) is indicated. B1a, B1b, and B2 cells from the peritoneal cavity (PeC) and B2 cells from the spleen (Spl) of control or Mb1-cre; *Whsc1*flSET mice were FACS sorted to over 95% purity. Each symbol represents one mouse. The experiment was done once with 2-3 mice per group. E) B1 cells do not develop in young Mb1-cre; *Whsc1*flSET (NSD2ΔSET) mice. FACS plots show relative abundance of B1a (IgM^hi^CD11b^+^CD5^+^) and B1b (IgM^hi^CD11b^+^ CD5^−^) cells in the peritoneal cavity of 3 week old mice. Absolute numbers of distinct lymphoid cells in the peritoneal cavity (bottom left) and percent (bottom right) are indicated. Representative plots of one experiment with 4-5 mice per group are shown.

Understanding the exact mechanisms of NSD2 contribution to B1 cell differentiation will require the development of approaches that allow for efficient inactivation of NSD2 in B1 cells during embryonic development as well as after establishment of the mature B1 cell compartment (maintenance question). However, already at this point our highly unexpected findings revealed NSD2 as the first-in-class epigenetic master regulator of a major B cell compartment in mice. This observation, while currently limited to studies in mice, may help to understand the contribution of NSD2 malfunction to lymphoid tumor development in human.

## Material and Methods

### Generation of *Whsc1*flSET mice

To create the targeting vector pBSmmsetflox, a single *loxP* site and a *BsoB*I restriction site and a Neo^r^ selection marker cassette flanked by FRT-sites were introduced into a HindIII site in intron 19 and an additional *loxP* site was inserted in a BglI site in intron 17 of the *Whsc1* locus. ES cells at embryonic day 14.1 were transfected and selected as described. Homologous recombinants were identified by Southern blot analysis (*BsoB*I digest 5’ probe or 3’ probe; *Nco*I digest 3’ probe). Targeted ES cells were used to generate mice. The FRT-site flanked Neo^r^ cassette was removed from targeted mice by breeding to Flip^r^ mice.

### Mice

*Whsc1*flSET mice were generated in our laboratory and are on a C57/Bl6 background. *Whsc1*flSET littermates without Cre were used as controls. Mice were housed under specific pathogen-free conditions and experimental protocols were approved by the Rockefeller University Institutional Animal Care and Use Committee. All studies were conducted in accordance with the GSK Policy on the Care, Welfare and Treatment of Laboratory Animals and were reviewed the Institutional Animal Care and Use Committee either at GSK or by the ethical review process at the institution where the work was performed.

### Antibodies

The following antibodies were purchased from either BD PharMingen, eBioscience or Jackson Labs: B220 (RA3-6B2), IA^b^ (AF6-120.1), IgM (115-116-075), IgD (11-26c.2a), CD5 (53-7-3), CD11b (M1/70), CD21/35 (7G6), CD19 (B3B4), CD23 (1D3), CD43 (S7), CD86 (GL1), CD90 (53-2.1). The B cell receptor specific antibodies PE-3H7 (anti-VH11id), and APC-13B5 (anti-Vk9id) were kindly provided by Kyoko Hayakawa (Fox Chase Cancer Center) and the NSD2 antibody (29D1) was purchased from Abcam.

### Histone post translational modification analysis

Histones were extracted in acid and chemically derivatized twice, digested with trypsin, followed two more rounds of derivatization and the peptides were desalted by using C_18_ stage-tips, as described earlier (Bhanu et al., 2015). Samples were analyzed using an EASY-nLC nanoHPLC (Thermo Scientific, Odense, Denmark) in a gradient of 0-35% solvent B (A = 0.1% formic acid; B = 95% MeCN, 0.1% formic acid) over 30 min and from 34% to 100% solvent B in 20 minutes at a flow-rate of 250 nL/min. Nano-liquid chromatography was coupled with a Q-Exactive mass spectrometer (Thermo Fisher Scientific, Bremen, Germany). Full scan MS spectrum (m/z 290−1650) was performed in the Orbitrap with a resolution of 30,000 (at 400 *m/z*) with an AGC target of 1×10e6. The MS/MS events included both data-dependent acquisition and target, the latter for isobaric peptides to enable MS/MS-based quantification. The relative abundance of histone H3 and H4 peptides were calculated by using EpiProfile.

### Flow cytometry

Single-cell suspensions from tissues were prepared. All antibodies were from BD Pharmingen, eBioscience, or Jackson Laboratories and were used at dilutions ranging from 1:100-1:3000 and incubated for 30 min at 4°C. Flow cytometric analysis and cell sorting were performed using a FACS LSR II or Aria (Becton Dickinson) and data were analyzed with FlowJo software (TreeStar).

### B-cell purification and *in vitro* proliferation

Splenic B cells were purified by depleting CD43^+^ cells, using anti-CD43 beads and magnetic columns (Milteyni Biotec) and stimulated *in vitro* with 10 μgml^−^ F(ab)2 fragment of goat anti-mouse IgM (Jackson ImmunoResearch) in combination with 25U ml^−^ recombinant mouse IL-4 (R&D), 5μgml^−^ bacterial LPS (Sigma), or 5 μgml^−^ bacterial LPS in combination with 25 Uml^−^ recombinant mouse IL-4 (R&D). Labeling of cells with 5-(and 6-) carboxyfluorescein diacetate, succinimidyl ester (CFDA-SE; Molecular Probes) for analysis of proliferation was performed following the manufacturer’s instructions. The decline in CFSE fluorescence as a measure of B-cell proliferation was determined by FACS analysis.

### Cell survival assay

Purified B cells were cultured either in medium alone or in the presence of 1.56-25ng ml^−^ of recombinant BAFF for the indicated time and stained with Annexin V (Roche) and 7-aminoactinomycin D (7-AAD; Sigma-Aldrich).

**Statistical analysis** was performed in Prism (Graphpad Software) with the unpaired t test for total cell numbers and frequencies. *P<0.05; **P<0.01; ***P<0.001

